# Biosensor optimization using a FRET pair based on mScarlet red fluorescent protein and an mScarlet-derived green fluorescent protein

**DOI:** 10.1101/2022.06.20.496847

**Authors:** Khyati Gohil, Sheng-Yi Wu, Kei Takahashi-Yamashiro, Yi Shen, Robert E. Campbell

## Abstract

Genetically encoded biosensors based on Förster resonance energy transfer (FRET) are indispensable tools for monitoring biochemical changes in cells. Green and red fluorescent protein-based FRET pairs offer advantages over the classically employed cyan and yellow fluorescent protein pairs, such as better spectral separation, lower phototoxicity, and less autofluorescence. Here, we describe the development of an mScarlet-derived green fluorescent protein (designated as mWatermelon) and its use as a FRET donor to the red fluorescent protein mScarlet-I as a FRET acceptor. We tested the functionality of this FRET pair by engineering biosensors for the detection of protease activity, Ca^2+^, and K^+^. Furthermore, we described a strategy to enhance the FRET efficiency of these biosensors by modulating the intramolecular association between mWatermelon and mScarlet-I.

## Introduction

Understanding biological processes requires tools that enable monitoring of dynamic changes in concentration or localization of various ions and molecules in the heterogeneous cellular environment. Fluorescent proteins (FPs), either on their own or as components of biosensors, are indispensable tools to probe these dynamic biochemical changes.^1^ A biosensor for an analyte or a cellular event typically consists of a recognition domain, which recognizes the binding of an analyte or occurrence of an event, and an indicator domain, which subsequently outputs a measurable signal.^2,3^ Fluorescent proteins are popular indicator domains for biosensor development as they generate a fluorescent signal which can be modulated in terms of intensity or wavelength. Coupled with microscopy technology, FP-based biosensors enable quantitative measurement and real-time imaging of biomolecules and cellular processes with high spatiotemporal resolution.

The availability of different hues of FPs has provided opportunities for constructing various genetically encodable biosensors. These biosensors can be categorized into two classes based on their response mechanism. One class consists of intensiometric biosensors in which a conformational change in the recognition domain is relayed to the fused single FP, leading to a modulation of the fluorescence intensity.^4^ The other class is based on recognition domain-dependent modulation of Förster resonance energy transfer (FRET) between two FPs of different fluorescent colours, leading to a ratiometric change in fluorescence emission.^5^ Numerous FRET-based biosensors have been engineered for the purpose of studying various intracellular processes such as enzyme activity, ion dynamics, post-translational modifications, and protein-protein interactions.^1,6^ These FRET-based biosensors are powerful molecular tools as they allow quantitative and noninvasive live-cell imaging in various model systems.

Canonical FRET-based biosensors typically employ cyan fluorescent proteins (CFPs) and yellow fluorescent proteins (YFPs), which remain the most effective and most commonly used FRET pair to date. Despite their popularity, CFP-YFP pairs have certain drawbacks such as violet excitation-induced phototoxicity, autofluorescence, and substantial spectral cross-talk.^6,7^ To overcome these issues, alternative FRET pairs with GFPs as the donor and an orange or red fluorescent protein (OFP or RFP) as an acceptor have been explored. The GFP-RFP pairing leads to reduced phototoxicity and improves spectral separation. The most prominent advancements in green-red FRET pairs for ratiometric imaging came from the development of Clover-mRuby2, and the further improved version mClover3-mRuby3, which has better photostability compared to existing GFP-RFP and CFP-YFP FRET pairs.^7,8^

To optimize the performance of FRET-based biosensors, multiple strategies have previously been explored.^5^ Improving the photophysical properties of FPs in a FRET pair is one way of improving FRET efficiency. Another way of increasing FRET efficiency is to adjust the relative orientation, or distance between, the two FPs. Since FRET is dependent on orientation and distance, these two factors can be modulated to obtain higher FRET efficiency leading to a better dynamic range (Δ*R/R*_min_, where R is the ratio of FRET acceptor intensity to donor intensity).^5^ A higher dynamic range enables a more sensitive FRET-based measurement of subtle conformational changes. One way to adjust the distance between two FPs is to optimize the linkers that connect the FPs to the recognition domain. Most biosensor development involves a critical step of optimizing the length and the molecular composition of these linkers.^9^ A second approach is to change the topology by incorporating a circularly permuted fluorescent protein (cpFP) as an acceptor or donor. A cpFP is a protein for which the topology has been changed by genetically joining the original C- and N-termini and generating new C- and N-termini elsewhere in the protein.^10,11^ A third approach is to adjust how the two FPs physically interact with each other by increasing the association between the two FPs such that a higher *R_max_* can be achieved^12^. This would ideally require an association that is fine-tuned such that the FRET-based biosensor can still form high FRET and low FRET states, depending on the conformational changes brought about by the binding of an analyte.^13–16^ The association should not be so strong that the FPs of the FRET pair perpetually dimerized and lose their ability to transduce any analyte binding changes.

To expand the FP-based FRET pair repertoire and diversify the potential optimization strategies for creating FRET-based biosensors with larger dynamic ranges, we have engineered a new green fluorescent protein based on mScarlet-I.^17^ This new GFP, paired with mScarlet-I, enabled the development of a new green-red FP-based FRET pair. As mScarlet is derived from a dimeric precursor, this new FRET pair provides an opportunity to introduce mutations within the original dimeric interface to promote the formation of weakly heterodimeric complexes. In this way, the FRET efficiency can be further modulated and optimized to produce FRET-based biosensors with improved performance.

## Results and Discussion

### Engineering of a green fluorescent protein-based on mScarlet-I

A favourable FRET acceptor needs to be as bright as possible in order to give a strong sensitized emission signal. Currently, mScarlet is one of the brightest red FPs. mScarlet is engineered from mRed7, a rationally designed synthetic RFP template primarily based on mCherry.^17^ The mScarlet protein sequence is phylogenetically close to the DsRed family of red FPs (**Figure S1A**) and shares similar surface residues (**Figure S1B**), thus is potentially amenable to oligomeric interface engineering.^17,18^ A variant of mScarlet called mScarlet-I matures faster, and has a better brightness when expressed in cells.^17^ Both mScarlet and mScarlet-I have been used as red FRET acceptors. Given the enhanced maturation and cellular brightness of mScarlet-I, we selected it as the preferred FRET acceptor for the green-red FRET pair.^19,20^

We hypothesized that a blue-shifted version of mScarlet-I could be a high-efficiency FRET donor, as it could potentially interact with mScarlet-I through self-associating oligomeric interactions which could be introduced by re-introduction of interface residues of the oligomeric ancestor of mScarlet-I (**Figure S2**). To achieve this, we first engineered a green FP from mScarlet-I. The development started with the site-directed saturation mutagenesis of position M67, the first residue of the chromophore-forming tripeptide. Within this library, we identified a dim green FP with the mutation M67G. This mutation changes the chromophore-forming tripeptide from MYG to GYG. Many green and yellow FPs have GYG as their chromophore forming tripeptide.^21,22^ We then performed five rounds of directed evolution with random mutagenesis library construction and brightness screening. This resulted in a bright green FP designated as mWatermelon (**Figure 1A**). From the template mScarlet-I, mWatermelon has a total of 8 mutations (V8L/F12Y/N24D/Q43H/M67G/Y121H/K139R/F178C) and none of these mutations reside at the A-C dimer interface (**Figure 1B, Figure S3**). Among the accumulated mutations, F178C, Y121H and Q43H face towards the chromophore and are most likely to affect the chromophore maturation and environment (**Figure 1B**).

**Figure 1.**
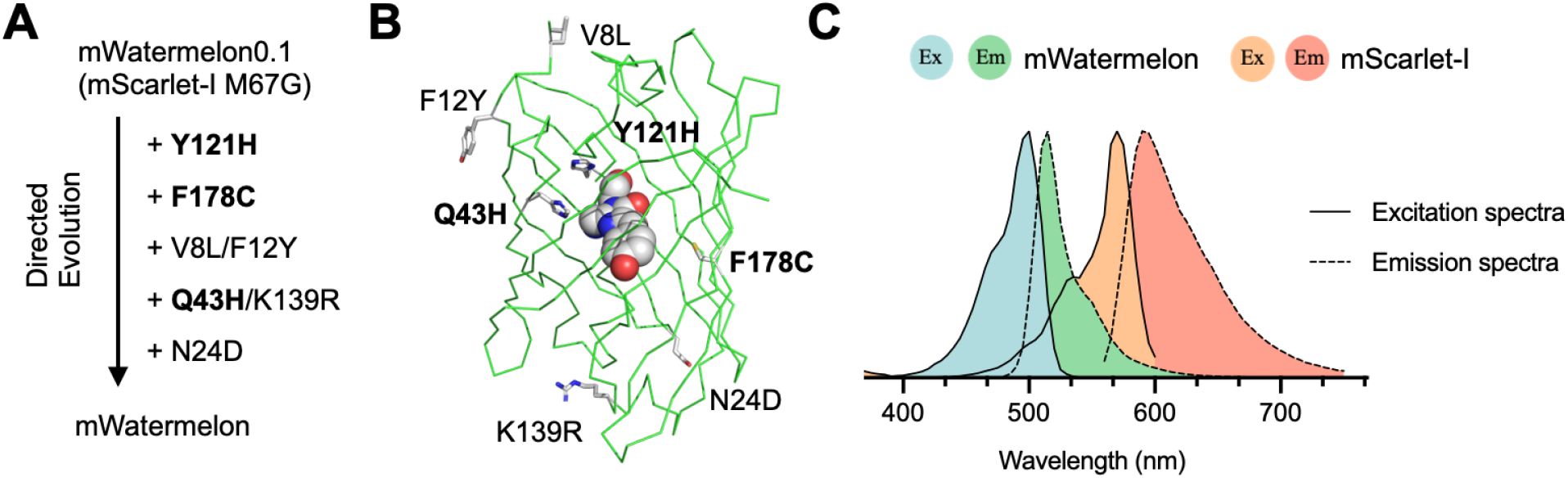
Development and properties of mWatermelon. (A) Evolution chart of mWatermelon. (B) Mutations mapped on mScarlet crystal structure (PDB: 5LK4). Mutations (gray sticks) highlighted in bold are facing towards the chromophore (sphere). (C) Excitation and emission spectra of mWatermelon and mScarlet-I.

This transformation of a red FP to a green FP is possible due to the proposed branched mechanism of chromophore formation in DsRed, which can lead to either a green chromophore species or a red chromophore species via a blue fluorescent intermediate.^23^ A previous example of the development of an FP with a hypsochromic shift in excitation and emission is mTagBFP, which was developed from TagRFP by structure-based directed evolution to trap the blue intermediate in the chromophore formation pathway and prevent its further maturation.^24,25^ Other examples are the development of ddGFP from ddRFP, and GGvT from tdTomato, which relied on two key mutations which were previously known to convert DsRed into a green FP.^12,26,27^

To characterize the key properties of mWatermelon, we measured the fluorescence spectrum, brightness, and pH sensitivity. The fluorescence excitation and emission maxima of mWatermelon are 504 nm and 515 nm, respectively (**Figure 1C**). The extinction coefficient of mWatermelon is 88,000 M^-1^cm^-1^ and the quantum yield is 0.47. A pH titration revealed that mWatermelon has a *pK_a_* of 5.2, which suggests that mWatermelon is relatively pH insensitive within the most physiologically relevant pH range (**Table 1**). Altogether, mWatermelon is a bright green fluorescent protein engineered from mScarlet-I. More importantly, with the unaltered dimeric interface from mScarlet-I, mWatermelon opens up the possibility of engineering weak dimerization with mScarlet-I for better FRET efficiency.

**Table 1.** Photophysical properties of mWatermelon compared to mScarlet-I.

|  | Extinction Coefficient<br>(EC, M<sup>-1</sup> cm<sup>-1</sup>) | Quantum Yield<br>(QY) | Molecular Brightness<br>(EC x QY, M<sup>-1</sup> cm<sup>-1</sup>) | Excitation Maxima<br>(nm) | Emission Maxima<br>(nm) | pK<sub>a</sub> |
| --- | --- | --- | --- | --- | --- | --- |
| mWatermelon | 88,000 | 0.47 | 41,000 | 504 | 515 | 5.2 |
| mScarlet-I | 104,000 | 0.54 | 56,000 | 569 | 593 | 5.4 |

### Green-red FP FRET-based protease biosensors

To assess the FRET efficiency between mWatermelon and mScarlet-I, we developed a protease biosensor based on mWatermelon and mScarlet-I. In FRET-based protease biosensors, the FRET donor and acceptor FPs are initially linked by a short protease substrate sequence and display high FRET efficiency. Following cleavage by protease, the dissociation of the two FPs leads to the complete loss of FRET. Therefore, a protease biosensor can serve as a convenient test system for benchmarking FRET pair performance (**Figure 2A**). To construct the protease sensor, mWatermelon and mScarlet-I were attached to a WELQut protease recognition peptide sequence (W-E-L-Q↓X) through a 16 amino acid flexible linker on either side of the recognition sequence (**Figure S4**). In an attempt to reduce the distance between the donor and the acceptor, we truncated the last six C-terminal residues on the N-terminal FP and the first four N-terminal residues on the C-terminal FP. First, the topological positioning of donor and acceptor FPs in the biosensor was assessed. The first topological arrangement tested had the donor mWatermelon on the N-terminus (biosensor named WPS, for m<u>W</u>atermelon-<u>p</u>rotease recognition peptide-m<u>S</u>carlet-I). The proteolysis of WPS resulted in a Δ*R/R_min_* of 2.56. Switching the positions of mWatermelon and mScarlet-I (biosensor named SPW) led to an increase in Δ*R/R_min_* to 3.18 (**Figure 2B**). This increase in FRET ratio change is mainly attributed to the higher FRET efficiency in the uncleaved state in SPW (**Figure 2C**). The improved FRET efficiency can be attributed to an increase in the percentage of mature mScarlet-I chromophores from 47.2% (WPS) to 57.1% (SPW), as estimated from the deconvolution of absorbance spectra (**Table S2** and **Figure S5**).

**Figure 2.**
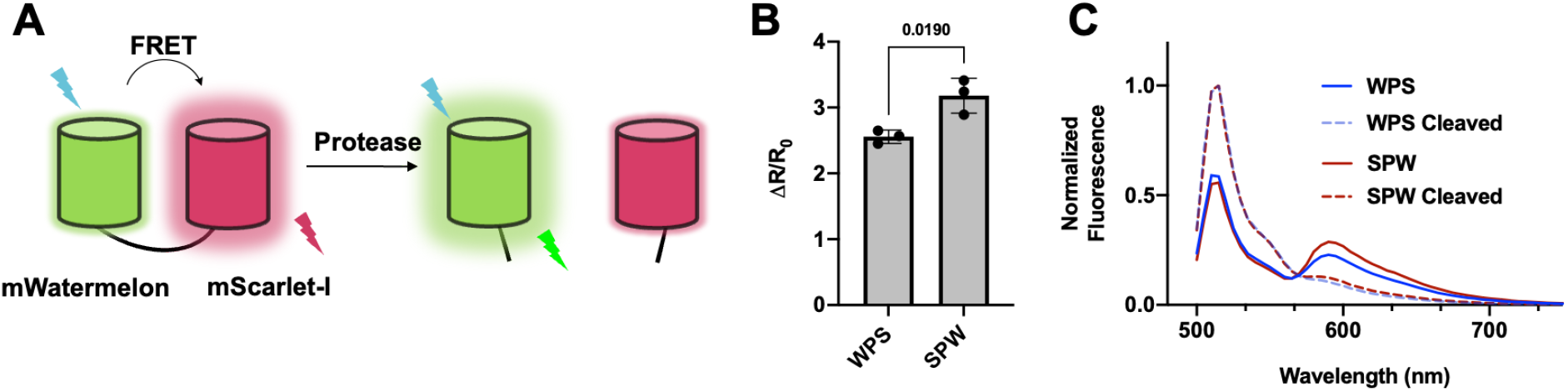
FRET-based protease biosensor. (A) Schematic representation of the biosensor constructs before and after being cleaved by WELQut protease. (B) FRET ratio change and (C) normalized emission spectra of protease biosensors mWatermelon-protease recognition peptide-mScarlet-I (WPS) and mScarlet-I-protease recognition peptide-mWatermelon (SPW). The bar graph shows mean ± S.D. for n = 3. The p-value is measured by an unpaired t-test and is indicated over the bar graph.

### Optimizing FRET efficiency by introducing dimeric interface mutations

To test if the intramolecular association strategy could be employed to further improve Δ*R/R*_min_, we focused on introducing weak dimeric interactions within the A-C dimer interface by reversing the mutations originally made in DsRed to break this interface and monomerize DsRed (**Figure S2**).^18,28^ Eleven mutations were introduced in order to break the A-C dimer interface of DsRed. Nine of these mutations are also present in mScarlet-I and two of the residues (A165, L175; residues numbered as found in mScarlet-I) match DsRed. We selected seven residues to reverse: E154R, K163H, R173H, A193Y, N195Y, S223H and T224L (**Figure 3A**). We systematically introduced these reversions by site-directed mutagenesis on both mWatermelon and mScarlet-I followed by measuring the change in the FRET dynamic range. The K163H mutation was made in all variants due to its central positioning at the A-C interface. The construct with all the seven mutations included on both FPs lead to a fully dimeric mScarlet-I and mWatermelon protease biosensor, designated as “dimeric SPW” (**Table S1**).

**Figure 3.**
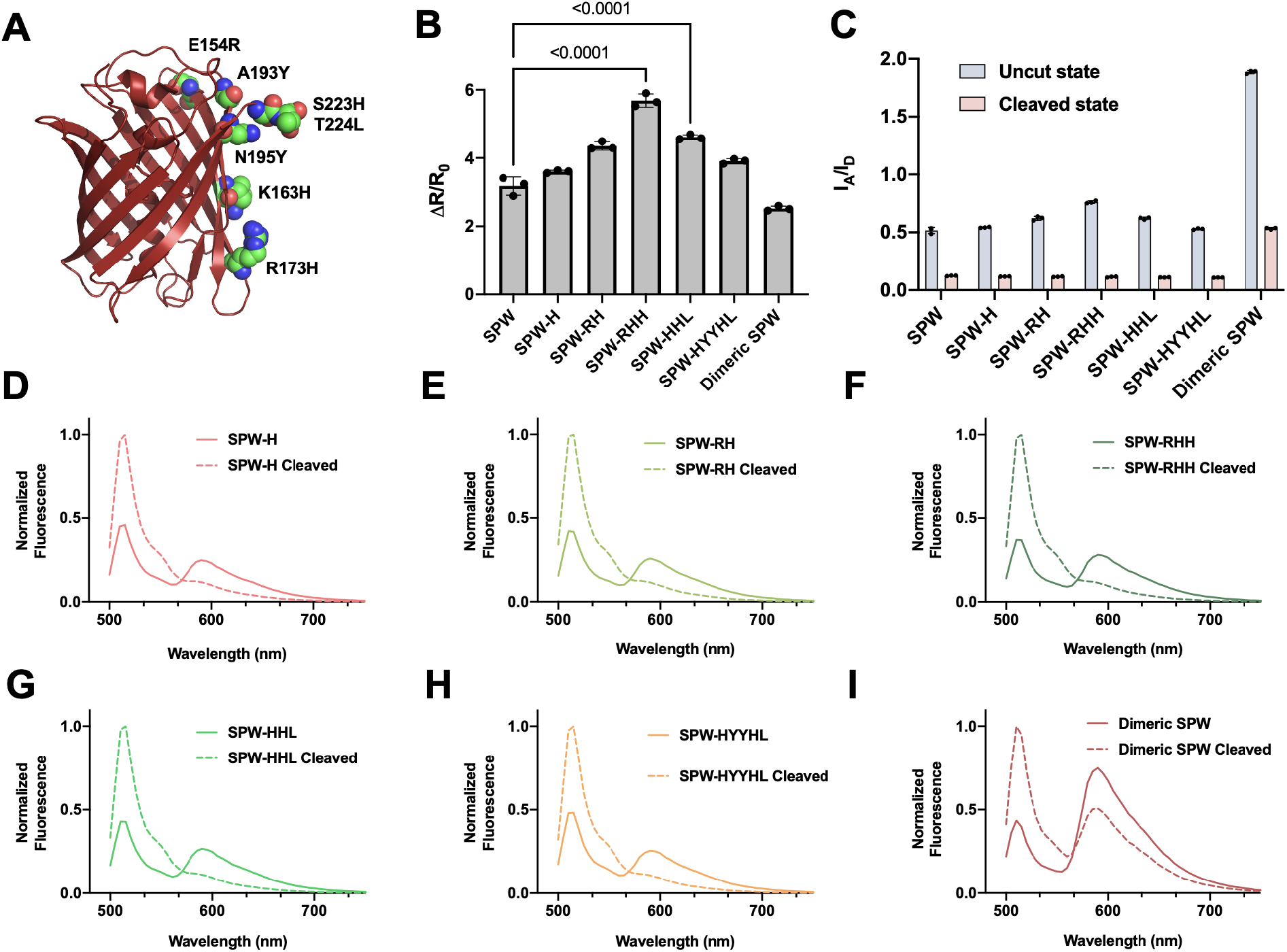
Optimizing FRET efficiency by introducing dimeric interface mutations. (A) Structure of mScarlet (PDB ID: 5LK4)^37^ showing key residues reversed on the interface in sphere representation. (B) Ratio changes and (C) FRET ratios before and after proteolysis of mScarlet-I-protease recognition peptide-mWatermelon (SPW) and variants with various combinations of mutations. The bar graph shows mean ± S.D. for n = 3. The p-value was measured by one-way ANOVA followed by Dunnett’s multiple comparison test vs. SPW and was indicated over the bar graph. Normalized emission spectra (n = 3) before and after cleavage of protease biosensors: (D) SPW, (E) SPW-RH, (F) SPW-RHH, (G) SPW-HHL, (H) SPW-HYYHL, and (I) dimeric SPW.

We characterized the FRET changes of these pairs in the context of the protease biosensor. In all variants, the FRET ratio (acceptor fluorescence intensity divided by donor fluorescence intensity, I_A_/I_D_) was higher in the uncleaved state as compared to the original SPW (**Figure 3B**), which is an indication that promoting intramolecular interactions helps to further improve the FRET efficiency. The FRET ratio in the cleaved state is expected to be the same for all variants as it depends primarily on the emission peak shape of the donor, and it was indeed observed to be the same for all variants except dimeric SPW. The dimeric SPW biosensor has a substantially higher FRET ratio than the original SPW in the cleaved state, which is attributable to a high-affinity interaction between dimeric mScarlet-I and dimeric mWatermelon preventing their dissociation (**Figure 3C**). This high FRET efficiency in the cleaved state of dimeric SPW leads to a reduced overall dynamic range. Other than dimeric SPW, all variants with different reversions exhibit higher ratio changes than the original SPW. The variants with 3 reversion mutations SPW-RHH (568 ± 19%) and SPW-HHL (461 ± 7%) have the highest FRET dynamic ranges, compared to 318 ± 26% for the original SPW (**Figure 3B**). The increase in FRET efficiency for most variants can be attributed to the increase in FRET ratio in the bound state (**Figure 3D-I**). Overall, the data suggested that mWatermelon is compatible with efforts to engineer the dimerization interface with mScarlet-I to increase their intramolecular association, and this strategy successfully improves the FRET dynamic range in FRET-based protease biosensors.

### Engineering of mWatermelon-mScarlet-I-based Ca^2+^ biosensors

Given the improved dynamic range of mWatermelon-mScarlet-I-based protease biosensors, we next expanded this strategy to develop biosensors for calcium ion (Ca^2+^). Ca^2+^ is an important and multi-functional secondary messenger involved in regulating various cellular processes such as growth, proliferation, and transcription.^29,30^ Due to the importance of Ca^2+^, many FP-based biosensors have been developed to study its dynamics.^31–33^ A FRET-based protease biosensor and a FRET-based Ca^2+^ biosensor function through different mechanisms. In protease biosensors, the irreversible cleavage of the substrate holding the FPs together results in the FPs becoming disconnected and freely diffusing apart. In the case of Ca^2+^ biosensors, the two FPs are linked together in both the absence and presence of Ca^2+^.

We constructed a FRET Ca^2+^ biosensor with mWatermelon-mScarlet-I pair using the canonical design of fusing the recognition domain consisting of calmodulin (CaM) and the CaM-binding peptide of myosin light-chain kinase (RS20) in between the two FPs (**Figure 4A, Figure S4**).^34^ When Ca^2+^ binds to CaM, an intramolecular interaction is initiated, which changes the CaM-RS20 conformation from extended to compact. In the Ca^2+^-bound compact state, the FRET ratio between mScarlet-I and mWatermelon showed an expected increase (**Figure 4B and 4C**). This Ca^2+^-induced response is reversible. Upon a decrease in Ca^2+^ concentration or chelation of Ca^2+^ by an external additive, the recognition domain should revert to its extended state, causing a decrease in FRET response (**Figure 4B**).

**Figure 4.**
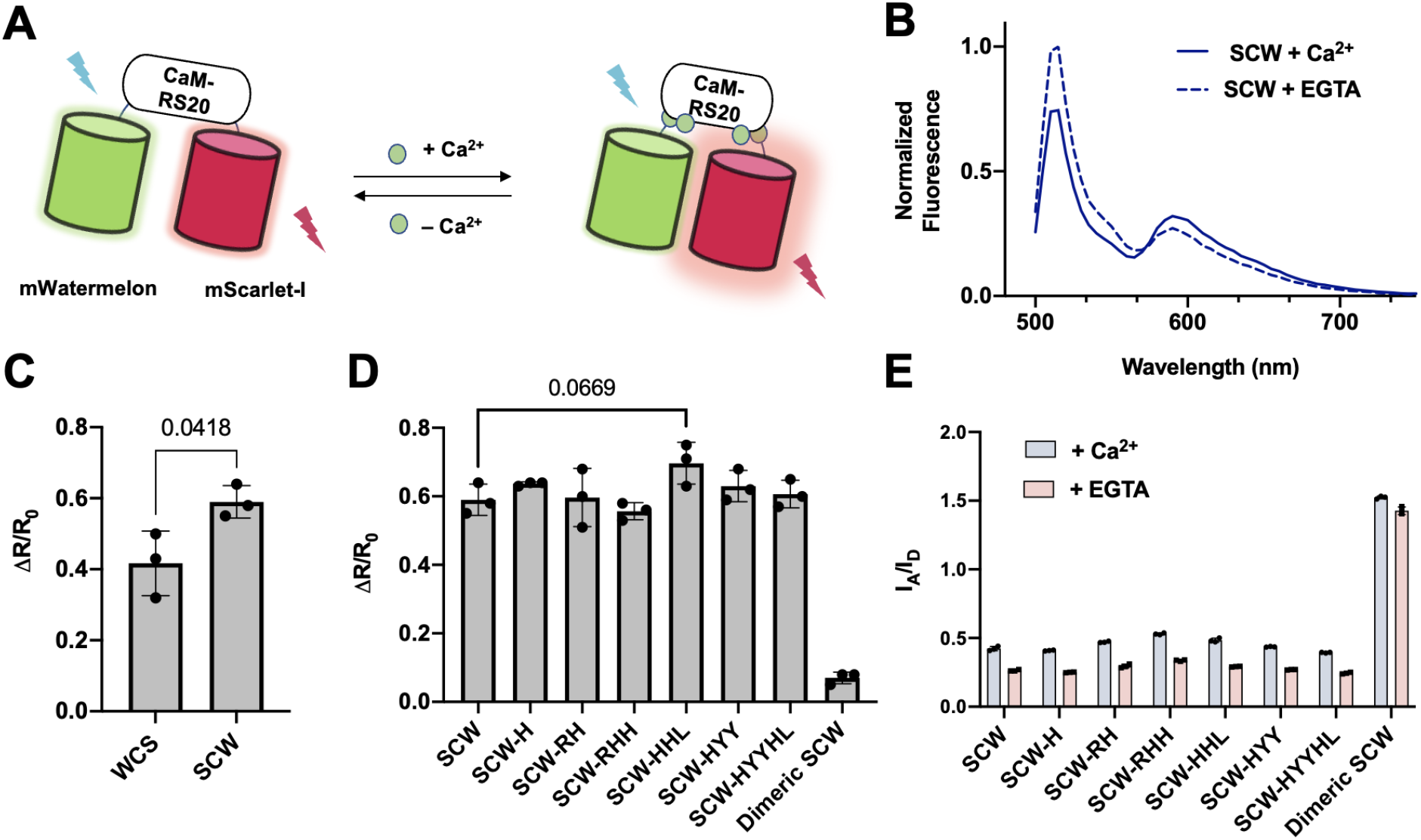
Design and characterization of mWatermelon-mScarlet-I-based Ca^2+^ biosensors. (A) Schematic representation of the mechanism of a mWatermelon-mScarlet-I-based Ca^2+^ biosensor. (B) Normalized fluorescence emission spectra of mScarlet-I-CaM-RS20-mWatermelon (SCW) with added Ca^2+^ and EGTA. (C) FRET ratio changes of Ca^2+^ biosensors mWatermelon-CaM-RS20-mScarlet-I (WCS) and SCW. (D) Ca^2+^ binding induced FRET ratio changes and (E) FRET ratios of SCW biosensor variants. Bar graphs show mean ± S.D. for n = 3. The p-values were measured by unpaired t-test for (C), and one-way ANOVA followed by Dunnett’s multiple comparison test for (D) vs. SCW, and relevant p-values are indicated over the bar graphs.

Following a strategy similar to the protease biosensor optimization, mutations that would increase the intramolecular association between the FPs were introduced with the aim of increasing the FRET dynamic range. We first assessed which arrangement of mScarlet-I and mWatermelon on the N- or C-terminus of the biosensor provided a better response. These biosensors are labelled as SCW or WCS based on the positioning of the FP (S = mScarlet-I, C = CaM-RS20 and W = mWatermelon). Having mScarlet-I on the N-terminus increased FRET efficiency by 17 ± 5 % (**Figure 4C**), which is likely a result of an increased percentage of mScarlet chromophores from 47.5% (WCS) to 56.9% (SCW) (**Table S2**). With the favoured FP arrangement established, the mutations E154R, K163H, R173H, A193Y, N195Y, S223H and T224L were again introduced systematically on both mWatermelon and mScarlet-I followed by measuring the change in dynamic range. When all these mutations were included on both FPs, the corresponding indicator is labelled as “dimeric SCW”. For most variants, FRET ratio (I_A_/I_D_) increases were observed in both Ca^2+^-rich and Ca^2+^-depleted states, leading to a small overall change in FRET ratios (**Figure 4D and 4E**). With the dimeric SCW, the two FPs seem to be tightly bound in both the presence and absence of Ca^2+^ due to the constitutive dimeric association of the FPs (**Figure 4E**), thus limiting its FRET dynamic range. Of all variants tested, a modest yet statistically insignificant (p = 0.067) increase was observed only with SCW-HHL (70% ± 6%) relative to SCW (59% ± 5%). Overall, we conclude that the dimerization interface engineering optimizing strategy was less effective on these Ca^2+^ biosensors compared to their protease-sensing counterparts, highlighting the differences in the biosensing mechanism and the need for further optimization.

### Improving green red FRET-based K^+^ biosensors with mWatermelon-mScarlet-I pair

The potassium ion (K^+^) is the most abundant intracellular cation and is critical for maintaining cellular functions. Dysregulation of K^+^ homeostasis can affect cardiovascular, neurological and the renal systems, leading to various diseases.^35^ The identification of *Escherichia coli* K^+^ binding protein (Kbp) led to the development of a series of biosensors, including the KIRIN1-GR green-red FRET-based K^+^ biosensor.^36^–^38^. Constructed with Clover-mRuby2, KIRIN1-GR has a rather limited *in vitro* FRET dynamic range *(ΔR/R_min_* of 20%).^38^ With the development of the mWatermelon-mScarlet-I FRET pair, we sought to engineer a new green-red K^+^ biosensor to further improve the FRET response. We thus developed such biosensors by fusing the mWatermelon-mScarlet FRET pair together with Kbp (**Figure 5A, 5B, Figure S4**). Gratifyingly, mScarlet-I-Kbp-mWatermelon (SKW) showed a substantial (~3x) increase in response over KIRIN1-GR (**Figure 5C**). SKW exhibits a *K_d_* of 1.31 ± 0.40 mM for K^+^, which is similar to the *K_d_* of KIRIN1-GR (2.56 ±0.01 mM for K^+^) (**Figure S6**). We then test if weak dimerizing mutants could further improve upon SKW. However, no further increase in FRET efficiency was observed in all variants with different reversions (**Figure 5D, 5E**). It is worth noting that in the variants of SKW-RHH and dimeric SKW, the FRET ratio (I_A_/I_D_) increases both in the presence of 150 mM K^+^ and the absence of K^+^, indicating the increased dimerization tendency resulted in lower FRET efficiency in the context of K^+^ biosensors. Although the dynamic range of SKW could not be improved further by the dimerization interface engineering strategy, this newly developed K^+^ biosensor provides a significantly better alternative to the original KIRIN1-GR.

**Figure 5.**
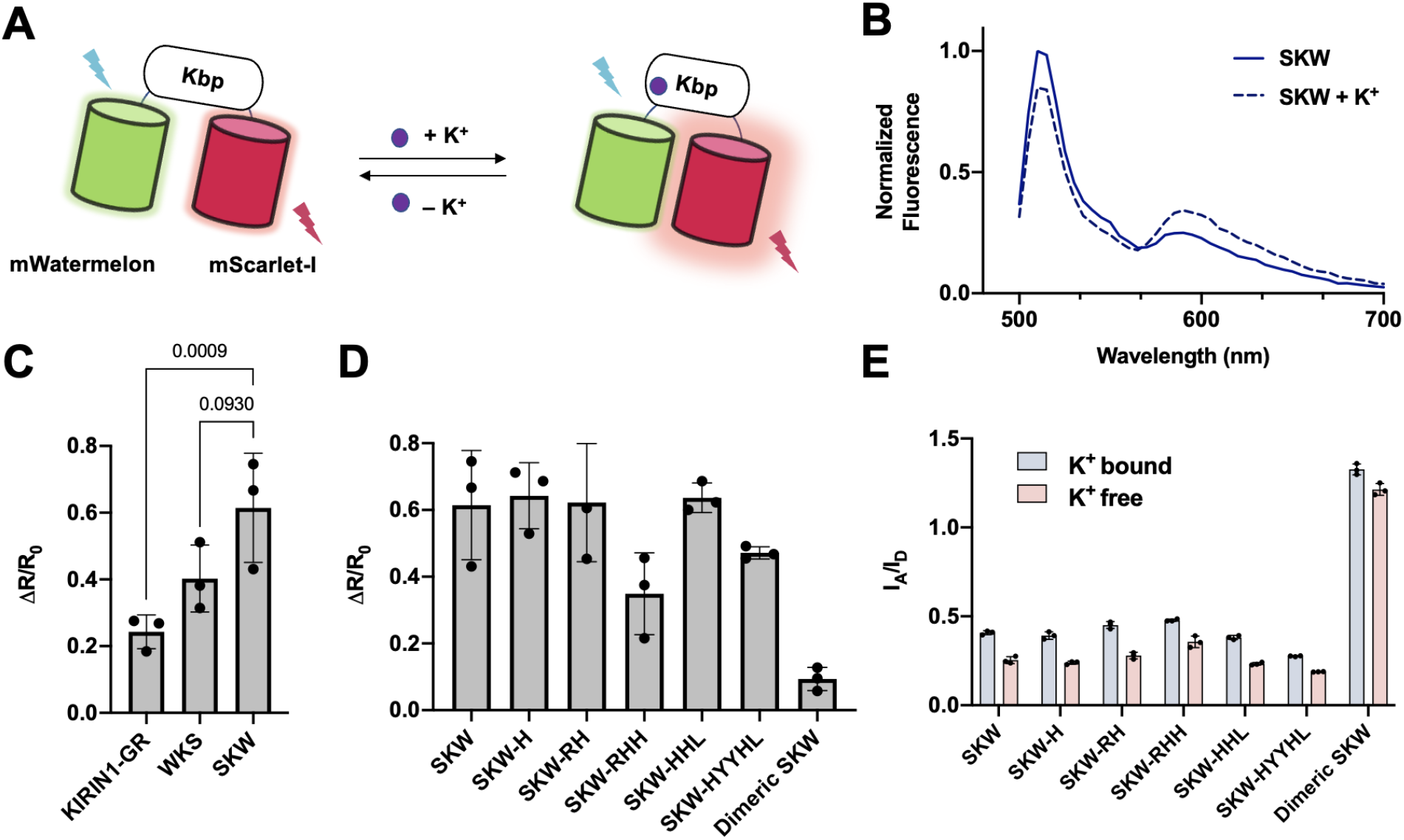
Design and characterization of mWatermelon-mScarlet-I-based K^+^ biosensors. (A) Schematic representation of the mechanism of FRET-based K^+^ biosensor. (B) Normalized emission spectra of mScarlet-Kbp-mWatermelon (SKW) in the absence and presence of 150 mM K^+^. (C) FRET ratio changes of K^+^ biosensors KIRIN1-GR, mWatermelon-Kbp-mScarlet (WKS), and SKW. (D) FRET ratio changes and (E) FRET ratios of SKW biosensor variants with mutations as listed on the x-axis. Data expressed as mean ± S.D. Bar graphs show mean ± S.D. for n = 3. The p-values were measured by an unpaired t-test for (C) and are indicated over the bar graph.

### Comparison of mWatermelon-mScarlet-I to other green-red FRET pairs

Having established that mWatermelon-mScarlet-I is a promising FP pair for developing green-red FRET biosensors for detecting protease activity, Ca^2+^, and K^+^, we next sought to compare its performance to the existing green-red FRET FP pairs. Towards this end, we selected the mClover3-mRuby3 (C-R) pair since it is one of the best available green-red FP pairs.^8^ More recently, ratiometric Ca^2+^ biosensors with green-red FPs were engineered and optimized using mScarlet-I as the FRET acceptor, along with GFP, Venus, mNeonGreen, and their circularly permuted variants as the FRET donors.^19^ Following extensive optimization, these biosensors exhibited high FRET efficiencies for imaging Ca^2+^ dynamics. We therefore also selected cpVenus175-mScarlet-I (cpV-S) pair from the highly responsive Twitch-VR Ca^2+^ biosensor.^20^ We then constructed protease, Ca^2+^, and K^+^ biosensors by substituting mWatermelon-mScarlet-I with these FRET pairs. For the sake of consistency, in all the biosensors the FRET donor was positioned on the N-terminus. mClover3-mRuby3 had the same number of residues deleted at the C-terminus and N-terminus, respectively, as mWatermelon-mScarlet-I.

In the protease biosensor group, mRuby3-mClover3 performed better than mScarlet-I-mWatermelon as a FRET pair, yet the interface-engineered SPW-RHH and SPW-HHL performed significantly better than both RPC and SPcpV (**Figure 6A**). For Ca^2+^ biosensors, SCW-HLL outperformed the mScarlet-I-CaM-RS20-cpVenus175 group and showed a similar response to mRuby3-CaM-RS20-mClover3 (**Figure 6B**). For the K^+^ biosensor, only a marginal decrease was observed in the biosensor with mRuby3-mClover3 as compared to mScarlet-I-mWatermelon and SKW-HHL (**Figure 6C**). Across all these three different biosensor groups, the mScarlet-I-mWatermelon HHL versions consistently performed equally well, or better, than their mScarlet-I-cpVenus and mRuby3-mClover3 counterparts (**Figure 6A-C**). In addition, photobleaching assays indicated comparable photostability for the mScarlet-I-mWatermelon pair compared to the other FRET pairs tested (**Figure S7, Table S3**). Overall, biosensors with the mScarlet-I-mWatermelon pairs offered comparable FRET responses to some of the current best green-red FRET pairs including mScarlet-I-cpVenus and mRuby3-mClover3. More importantly, the dimeric interface engineered variants mScarlet-I-mWatermelon HHL consistently performed well in the comparisons across different categories, suggesting it is an effective and reasonable starting point for the development of new green-red FRET-based biosensors.

**Figure 6.**
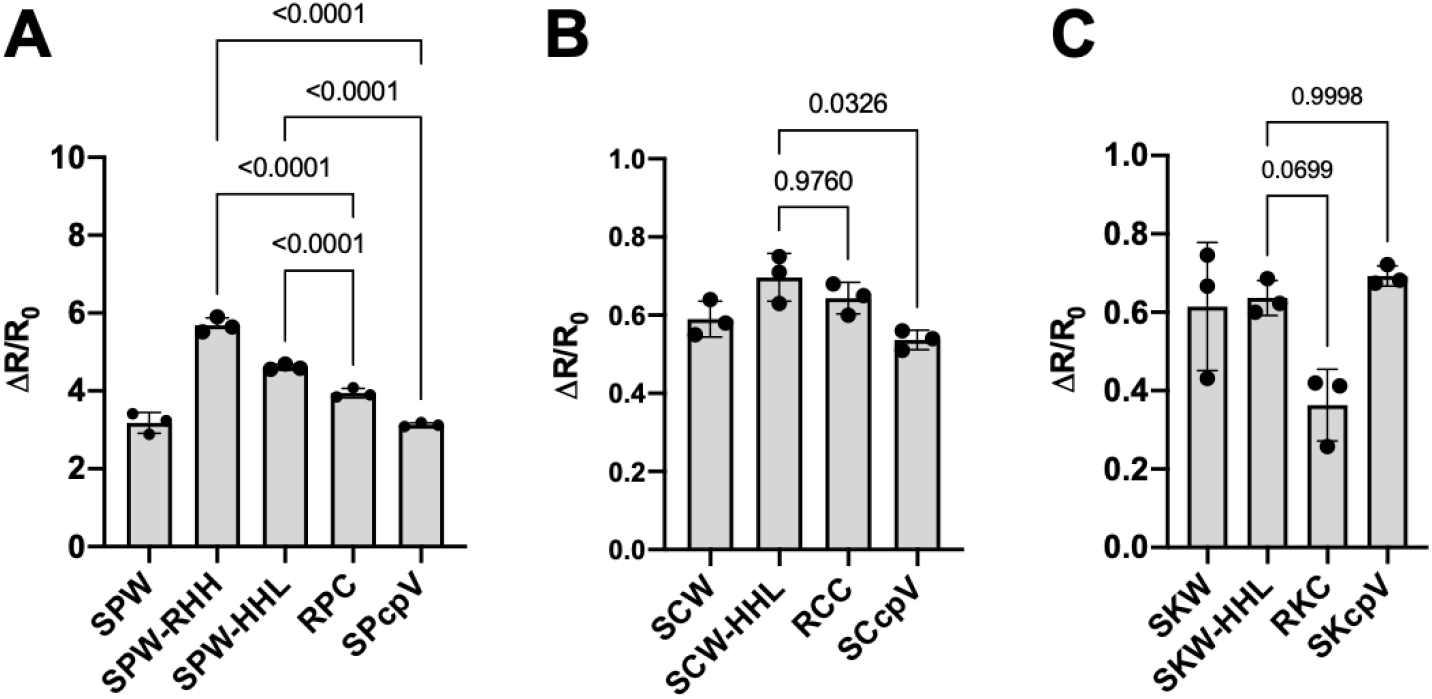
Benchmarking the performance of mWatermelon-mScarlet-I biosensors in vitro. FRET ratio changes of (A) protease biosensors (B) Ca^2+^ biosensors (C) K^+^ biosensors. Bar graphs show mean ± S.D. for n = 3. The p-values were measured by one-way ANOVA followed by Dunnett’s multiple comparison test, and are indicated over the bar graph.

### Fluorescence imaging of new Ca2+ biosensors in mammalian cells

To further evaluate the performance of the mWatermelon-mScarlet-I-based Ca^2+^ biosensor, we compared SCW and SCW-HHL, along with RCC and SCcpV in mammalian cells. We expressed and imaged the SCW, SCW-HHL, RCC and SCcpV biosensors in HeLa cells under identical experimental conditions. SCW, SCW-HHL, and SCcpV all exhibited similar ratios of fluorescence intensities in the green channel (ch1; excitation 473 nm, emission 515/25 nm) relative to the FRET channel (ch2; excitation 473 nm, emission 625/50 nm). In contrast, RCC exhibited a substantially dimmer signal in the FRET channel (**Figure 7A**), suggesting a lower FRET efficiency between mClover3 and Ruby3. We next assessed biosensor responses to histamine-induced intracellular Ca^2+^ oscillations (**Figure 7B**). The mWatermelon-mScarlet-I-based SCW and SCW-HHL, and mRuby3-mClover3-based RCC had similar responses in terms of maximum FRET change Δ*R*/*R*_0_, which was significantly better than the response of mScarlet-I-cpVenus175-based SCcpV (**Figure 7C**). In terms of signal-to-noise ratio (SNR), both SCW and SCW-HHL outperformed RCC and SCcpV (**Figure 7D**). These results suggest that the mWatermelon-mScarlet-I FRET pair works as well or better than existing FRET pairs in cultured mammalian cells.

**Figure 7.**
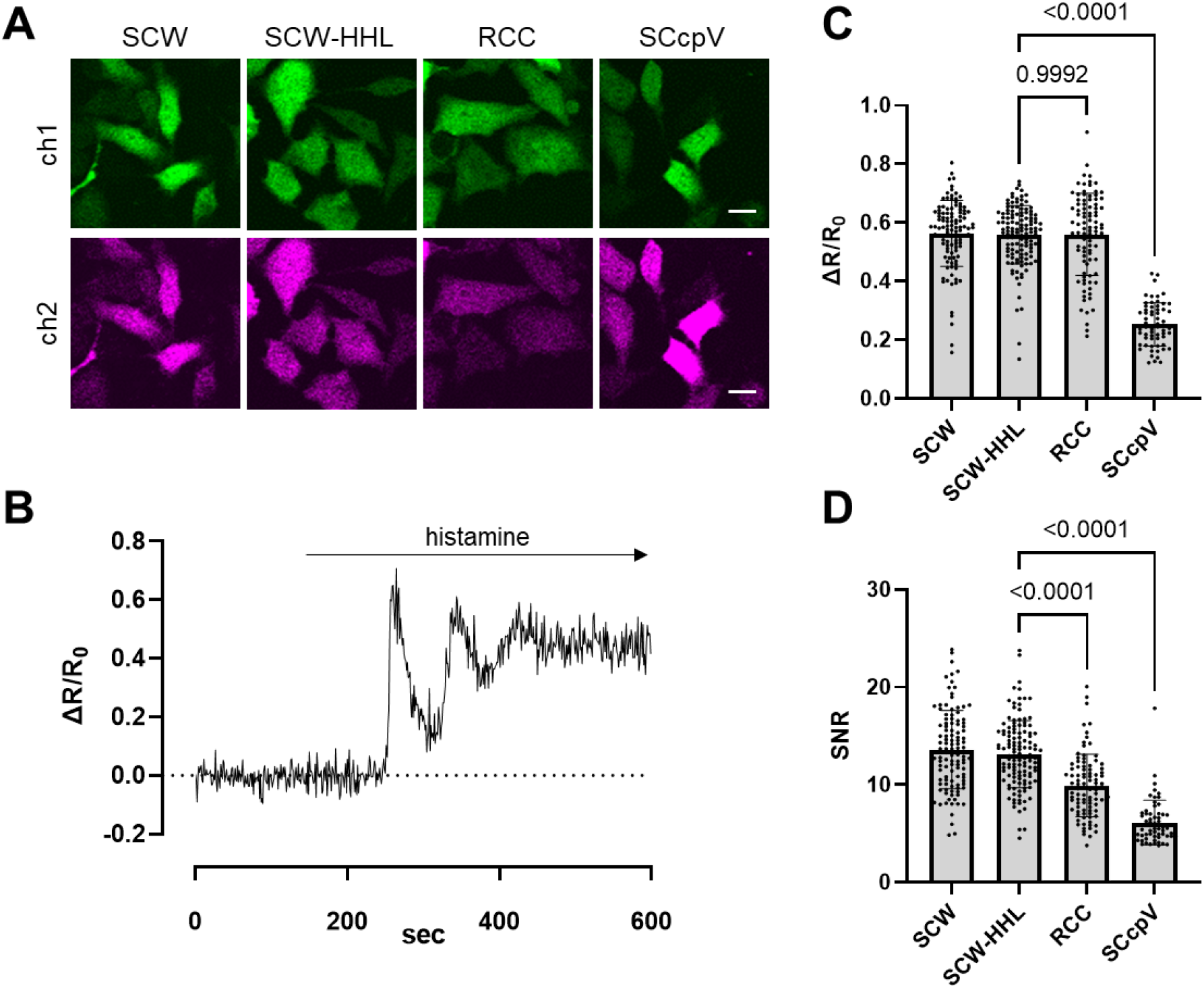
Performance of mWatermelon-mScarlet-I-based Ca^2+^ biosensors in HeLa cells. Ca^2+^ biosensors SCW, SCW-HHL, RCC and SCcpV were expressed in HeLa cells. (A) Representative fluorescence images of HeLa cells expressing Ca^2+^ biosensors in the cytosol (Scale bar = 20 μm) (excitation; 473 nm, emission; ch1 515/25 nm, ch2 625/50 nm). (B) Representative FRET ratio change of SCW-HHL Ca^2+^ biosensor with histamine stimulation. FRET ratio changes of Ca^2+^ biosensors (C) and their signal to noise ratio plots (D) (SCW n = 112 cells, SCW-HHL n = 127 cells, RCC n = 94 cells, SCcpV; n = 62 cells). The experiments were repeated more than three times independently and the combined results are shown. Bar graphs show mean ± S.D. The p-values were measured by one-way ANOVA followed by Dunnett’s multiple comparison test, and are indicated over the bar graph.

## Conclusions

Here we described the development of a new green-red FRET pair with mScarlet-I as the FRET acceptor and mWatermelon (a green fluorescent version of mScarlet-I) as the FRET donor. The potential of this FP pair has been established by developing FRET-based biosensors to detect protease activity, Ca^2+^, and K^+^. The development of mWatermelon enables a strategy to improve the dynamic range of mScarlet-I-mWatermelon biosensors by introducing hydrophobic mutations which would promote weak hetero-dimerization of mScarlet-I and mWatermelon. Although this strategy proved useful in improving FRET efficiency for our protease biosensor, it was less effective in improving the FRET efficiency of the Ca^2+^ and K^+^ biosensors. This suggests that the strategy can be useful when mScarlet-I and mWatermelon are closer in proximity and are joined together by flexible linkers that allow the orientational flexibility to form intramolecular interactions. When mScarlet-I and mWatermelon are fused along with a comparatively rigid binding domain, their conformational mobility is more limited, and they may not be able to adopt the orientation required for intramolecular interactions to occur. Constitutive high-affinity dimerization, as seen in dimeric SCW and dimeric SKW, locks the two FPs in a bound state, essentially rendering the biosensor unable to respond to any changes in the recognition domain. A finely balanced equilibrium is required to ensure that FP interactions are driven primarily by the change in conformation brought about by the binding domains and that any other enhancement to FP interaction is weak enough to improve FRET efficiency in the bound state but does not affect the unbound state.

Among all the dimerizing mutants, we identified a set of mutations HHL that could serve as a good starting point for green-red FRET biosensor development, across different biosensor types, with mWatermelon and mScarlet-I. We also strongly suggest that biosensors need to be optimized individually based on their own unique binding domains. Further optimization strategies include improving the photophysical properties of these FPs, linker optimization, systematically engineering topological variants, and further dimerization interface engineering to fine-tune the affinity. Overall, mScarlet-I and mWatermelon are a robust FP FRET pair that could be utilized to create FRET biosensors for various cellular applications.

## Materials and Methods

### General methods and material

All DNA primers for cloning and mutagenesis were purchased from Integrated DNA Technologies (IDT). Genes encoding individual components of biosensors were amplified by polymerase chain reaction (PCR) using Q5^®^ High-Fidelity DNA Polymerase (New England Biolabs). Site-directed mutagenesis was performed by QuikChange lightning mutagenesis kit (Agilent) or by CloneAmp HiFi PCR Premix (Takara Bio). Random mutagenesis was performed using Error-Prone PCR (EP-PCR) with Taq DNA polymerase (New England Biolabs). PCR products and restriction digestion products were purified using a GeneJET gel extraction kit (ThermoFisher). Restriction enzymes and ligases were purchased from New England Biolabs or ThermoFisher. Absorbance spectra were recorded with a DU-800 UV-visible spectrophotometer (Beckman). The fluorescence spectra were measured using a Safire2 microplate reader (Tecan).

### Protein engineering and biosensor construction

Engineering of mWatermelon was done via site-directed mutagenesis and multiple rounds of error-prone PCR (EP-PCR) using plasmids encoding mScarlet-I as template. Variant libraries were screened for brighter green fluorescence using a custom-built fluorescent colony imaging system. The initial protease biosensor was constructed by amplifying the gene encoding mWatermelon and mScarlet-I by PCR and overlapping these PCR products one at a time with a DNA oligomer that encodes for the synthesized linker and WELQ protease substrate. Ca^2+^ and K^+^ biosensors were constructed similarly. All other genes encoding the FP variants of mWatermelon, variants of mScarlet-I, cpVenus, mRuby3 and mClover3 were amplified by PCR and inserted into these initial constructs by restriction digestion followed by ligation of one FP gene at a time (restriction digestion sites shown in **Figure S5**). These constructed DNA encoding various biosensors were then assembled into a pBAD expression vector. All biosensor constructs were verified by DNA sequencing.

### Protein purification and *in vitro* characterization

The assembled pBAD plasmids were then used to transform *E. coli* strain DH10B by electroporation. The transformed DH10B cells were plated onto agar with LB medium containing 0.4 mg/ml ampicillin and 0.02% w/v L-arabinose and grown overnight at 37 °C. A single colony for each biosensor was picked and cultured in 150 mL of LB medium containing 0.1 mg/ml ampicillin and 0.02% w/v L-arabinose and incubated in a shaker (225 rpm) at 25 °C for 48 hours. Bacteria were harvested at 10,000 rpm, 4 °C for 10 min and the cell pellet was then resuspended in 25 mL of 1× Tris-buffered saline (TBS). Halt™ protease inhibitor cocktail (ThermoFisher) was added to the resuspended pellet and the cells were then lysed using sonication. The lysate was clarified by centrifugation at 10,000 rpm for 30 min. All the biosensor proteins have an N-terminal His tag, which allows for their purification from the supernatant by affinity chromatography using Ni-NTA agarose resin. The eluted protein solution was then buffer exchanged using a PD-10 (GE Healthcare Life Sciences) desalting column.

### *In vitro* characterization

Extinction coefficient of purified mWatermelon protein was determined by the alkali denaturation method. The absorption spectrum was measured with and without 1M NaOH. The extinction coefficient was calculated with the assumption that in 1M NaOH solution GFP-type chromophores have an extinction coefficient of 44,000 M^-1^cm^-1^ at their absorption peaks near 450 nm. The quantum yield of purified mWatermelon protein was determined by using mGreenLantern as a standard. The emission fluorescence spectrum of a dilution series of mWatermelon/mGreenLantern (absorbance corresponding to 0.01 to 0.1) was measured. The emission fluorescence was integrated and plotted against the absorbance. The slope from this plot was used to calculate the quantum yield of mWatermelon. The relative amounts of mWatermelon and mScarlet-I present in various biosensors were calculated from the measured absorbance spectra of the biosensors. First, mWatermelon and mScarlet-I spectra were deconvoluted from the biosensor’s absorbance spectra. Then the relative amount of FP was calculated by dividing the peak absorbance by the EC of the FP.

Fluorescence spectra of protease, Ca^2+^, and K^+^ biosensors were measured with a Tecan Safire2 microplate reader. The protease biosensor proteins (approximately 25 μg) were incubated with WELQut protease (final concentration of 0.1 unit in TBS) at 25 °C for 12 hours. The emission spectrum was measured for both uncleaved and proteolyzed biosensors. The fluorescence spectrum for Ca^2+^ biosensor proteins was measured in the presence of 39 μM free Ca^2+^ (10 mM CaEGTA in 100 mM KCl, 30 mM MOPS, pH 7.2) and zero free Ca^2+^ buffer (10 mM EGTA in 100 mM KCl, 30 mM MOPS, pH 7.2). The fluorescence spectrum for K^+^ biosensor proteins was measured in the presence and absence of 150 mM K^+^. For K^+^ *K_d_* determination, the purified protein was diluted into a series of buffers with K^+^ concentrations ranging from 0 to 150 mM. The biosensors that have mWatermelon or mClover3 were excited at 470 nm and the emission spectra were scanned from 500 to 750 nm whereas the biosensors that had cpVenus were excited at 480 nm and emission spectra were scanned from 510 to 750 nm.

Photobleaching kinetics assays were performed on an Axiovert 200 M (Zeiss) microscope equipped with a 75-W xenon-arc lamp, 40× objective lens (NA = 1.3, oil), and a digital CMOS camera (Orca Flash 40; Hamamatsu). The protein samples were vortexed with mineral oil in an Eppendorf tube and prepared on a coverslip as droplets in mineral oil. The protein droplets were under continuous illumination with a 565/50 nm excitation filter (Semrock) for red FPs (mScarlet-I, mScarlet-HHL, and mRuby) and a 488/50 nm excitation filter (Semrock) for green/yellow FPs (mWatermelon, mWatermelon-HHL, mClover, and cpVenus175). The images of these droplets (n ≥ 3) were acquired with Metamorph software (Molecular Devices) with a 30-s interval and a 100-ms exposure time.

### Live cell imaging

Live-cell imaging was performed with HeLa cells, which were obtained from the European Collection of Authenticated Cell cultures (ECACC). Cells were cultured in Dulbecco’s Modified Eagle’s Medium (DMEM, Gibco) supplied with 10% fetal bovine albumin serum. Cells were seeded on 12-mm coverslips coated with poly-L-lysine (MilliporeSigma) in 12-well plates. Cells were transfected with biosensor encoding plasmids using lipofectamine 3000 (ThermoFisher). Live-cell imaging was performed 48 hours post-transfection with a laser-scanning confocal microscope (FV1000, Olympus) equipped with a 20× objective lens (XLUMPLFLN, NA = 1.0, water). The laser and wavelength settings are as follows: ch1; excitation 473 nm and emission 515/25 nm, ch2 (FRET); excitation 473 nm and emission 625/50 nm. For imaging, the coverslips were transferred to the chamber filled with Hanks’ Balanced Salt Solution (HBSS) containing Ca^2+^ and Mg^2+^ (Gibco). Cells were treated with histamine (50 μM, MilliporeSigma) using a perfusion system. The flow rate was 5 ml/min via a peristaltic pump (Watson-Marlow Alitea-AB; Sin-Can). Fluorescence images were analyzed by Fiji ImageJ software. Time Series Analyzer plugin was used for the time-dependent quantification. For calculating the responses, non-responding samples were discarded.

### Statistical analysis

All data are expressed as individual data points or mean ± S.D. Sample sizes (n) are listed for each experiment. Statistical analyses and graphing were done using GraphPad Prism version 9.2.0. p-values were determined by one-way ANOVA followed by Dunnett’s multiple comparisons test or unpaired t-tests as mentioned in figure captions. *K*d curves were simulated by GraphPad Prism using a dose-response curve model with variable slope (four parameters) with the least-squares fit.

## Author Contributions

KG assembled DNA constructs, performed *in vitro* characterization, analyzed data, prepared figures, and wrote the manuscript. SYW performed *in vitro* characterization, analyzed data, prepared figures, and wrote the manuscript. KTY performed live cell imaging, analyzed data and prepared figures. YS performed directed evolution and *in vitro* characterization, supervised research, analyzed data, prepared figures, and wrote the manuscript. REC supervised research. All authors contributed to the editing and proofreading of the manuscript.

## Data and code availability

Data supporting the findings in this research are available from the corresponding author upon request. Code for spectral deconvolution and FP stoichiometry calculation is available at: https://github.com/shengyi2/spectrum_analysis/. Plasmids and DNA sequences will be available via Addgene.

## Acknowledgements

This work was supported by grants from the Canadian Institutes of Health Research (CIHR, FS-154310 to REC) and the Natural Sciences and Engineering Research Council of Canada (NSERC, RGPIN 2018-04364 to REC, RGPIN-2020-05514 to KB, and RGPIN-2016-06478 to MJL). SYW was supported by NSERC Canada Graduate Scholarships-Doctoral program, Alberta Innovates Technology Future (AITF) Graduate Scholarship, and the University of Alberta. KTY was supported by the Uehara memorial postdoctoral fellowship. We thank Dr. Simmen and Dr. Bassot for their support, Dr. Ballanyi and Dr. Rancic for technical assistance with a confocal microscopy. We also thank the University of Alberta Molecular Biology Services Unit (MBSU) for DNA sequencing support.

## Supplementary Information

**Supplementary Table 1.** FRET biosensors constructed in this study.

| FRET biosensor | N-terminal FP | Sensing domain | C-terminal FP |
| --- | --- | --- | --- |
| WPS | mWatermelon-ΔC6 | WELQut protease substrate | mScarlet-I-ΔN4 |
| SPW | mScarlet-I-ΔC6 | WELQut protease substrate | mWatermelon-ΔN4 |
| SPW-H | mScarlet-I-ΔC6 K163H | WELQut protease substrate | mWatermelon-ΔN4 K163H |
| SPW-RH | mScarlet-I-ΔC6<br>E154R/K163H | WELQut protease substrate | mWatermelon-ΔN4<br>E154R/K163H |
| SPW-RHH | mScarlet-I-ΔC6<br>E154R/K163H/R173H | WELQut protease substrate | mWatermelon-ΔN4<br>E154R/K163H/R173H |
| SPW-HHL | mScarlet-I-ΔC6<br>K163H/S223H/T224L | WELQut protease substrate | mWatermelon-ΔN4<br>K163H/S223H/T224L |
| SPW-HYYHL | mScarlet-I-ΔC6<br>K163H/A193Y/N195Y/S<br>223H/T224L | WELQut protease substrate | mWatermelon-ΔN4<br>K163H/A193Y/N195Y/S223H/T<br>224L |
| Dimeric SPW | mScarlet-I-ΔC6<br>E154R/K163H/R173H/A<br>193Y/N195Y/S223H/T2<br>24L | WELQut protease substrate | mWatermelon-ΔN4<br>E154R/K163H/R173H/A193Y/N<br>195Y/S223H/T224L |
| WCS | mWatermelon-ΔC6 | CaM-RS20 | mScarlet-I-ΔN4 |
| SCW | mScarlet-I-ΔC6 | CaM-RS20 | mWatermelon-ΔN4 |
| SCW-H | mScarlet-I-ΔC6 K163H | CaM-RS20 | mWatermelon-ΔN4 K163H |
| SCW-RH | mScarlet-I-ΔC6<br>E154R/K163H | CaM-RS20 | mWatermelon-ΔN4<br>E154R/K163H |
| SCW-RHH | mScarlet-I-ΔC6<br>E154R/K163H/R173H | CaM-RS20 | mWatermelon-ΔN4<br>E154R/K163H/R173H |
| SCW-HHL | mScarlet-I-ΔC6<br>K163H/S223H/T224L | CaM-RS20 | mWatermelon-ΔN4<br>K163H/S223H/T224L |
| SCW-HYY | mScarlet-I-ΔC6<br>K163H/A193Y/N195Y | CaM-RS20 | mWatermelon-ΔN4<br>K163H/A193Y/N195Y |
| SCW-HYYHL | mScarlet-I-ΔC6<br>K163H/A193Y/N195Y/S<br>223H/T224L | CaM-RS20 | mWatermelon-ΔN4<br>K163H/A193Y/N195Y/S223H/T<br>224L |
| Dimeric SCW | mScarlet-I-ΔC6<br>E154R/K163H/R173H/A<br>193Y/N195Y/S223H/T2<br>24L | CaM-RS20 | mWatermelon-ΔN4<br>E154R/K163H/R173H/A193Y/N<br>195Y/S223H/T224L |
| WKS | mWatermelon-ΔC6 | Kbp | mScarlet-I-ΔN4 |
| SKW | mScarlet-I-ΔC6 | Kbp | mWatermelon-ΔN4 |
| SKW-H | mScarlet-I-ΔC6 K163H | Kbp | mWatermelon-ΔN4 K163H |
| SKW-RH | mScarlet-I-ΔC6<br>E154R/K163H | Kbp | mWatermelon-ΔN4<br>E154R/K163H |
| SKW-RHH | mScarlet-I-ΔC6 | Kbp | mWatermelon-ΔN4 |
|  | E154R/K163H/R173H |  | E154R/K163H/R173H |
| SKW-HHL | mScarlet-I-ΔC6<br>K163H/S223H/T224L | Kbp | mWatermelon-ΔN4<br>K163H/S223H/T224L |
| SKW-HYYHL | mScarlet-I-ΔC6<br>K163H/A193Y/N195Y/S<br>223H/T224L | Kbp | mWatermelon-ΔN4<br>K163H/A193Y/N195Y/S223H/T<br>224L |
| Dimeric SKW | mScarlet-I-ΔC6<br>E154R/K163H/R173H/A<br>193Y/N195Y/S223H/T2<br>24L | Kbp | mWatermelon-ΔN4<br>E154R/K163H/R173H/A193Y/N<br>195Y/S223H/T224L |
| RPC | mRuby3-ΔC6 | WELQut protease substrate | mClover3-ΔN4 |
| RCC | mRuby3-ΔC6 | CaM-RS20 | mClover3-ΔN4 |
| RKC | mRuby3-ΔC6 | Kbp | mClover3-ΔN4 |
| SPcpV | mScarlet-I-ΔC6 | WELQut protease substrate | cpVenus175 |
| SCcpV | mScarlet-I-ΔC6 | CaM-RS20 | cpVenus175 |
| SKcpV | mScarlet-I-ΔC6 | Kbp | cpVenus175 |

**Supplementary Table 2.**
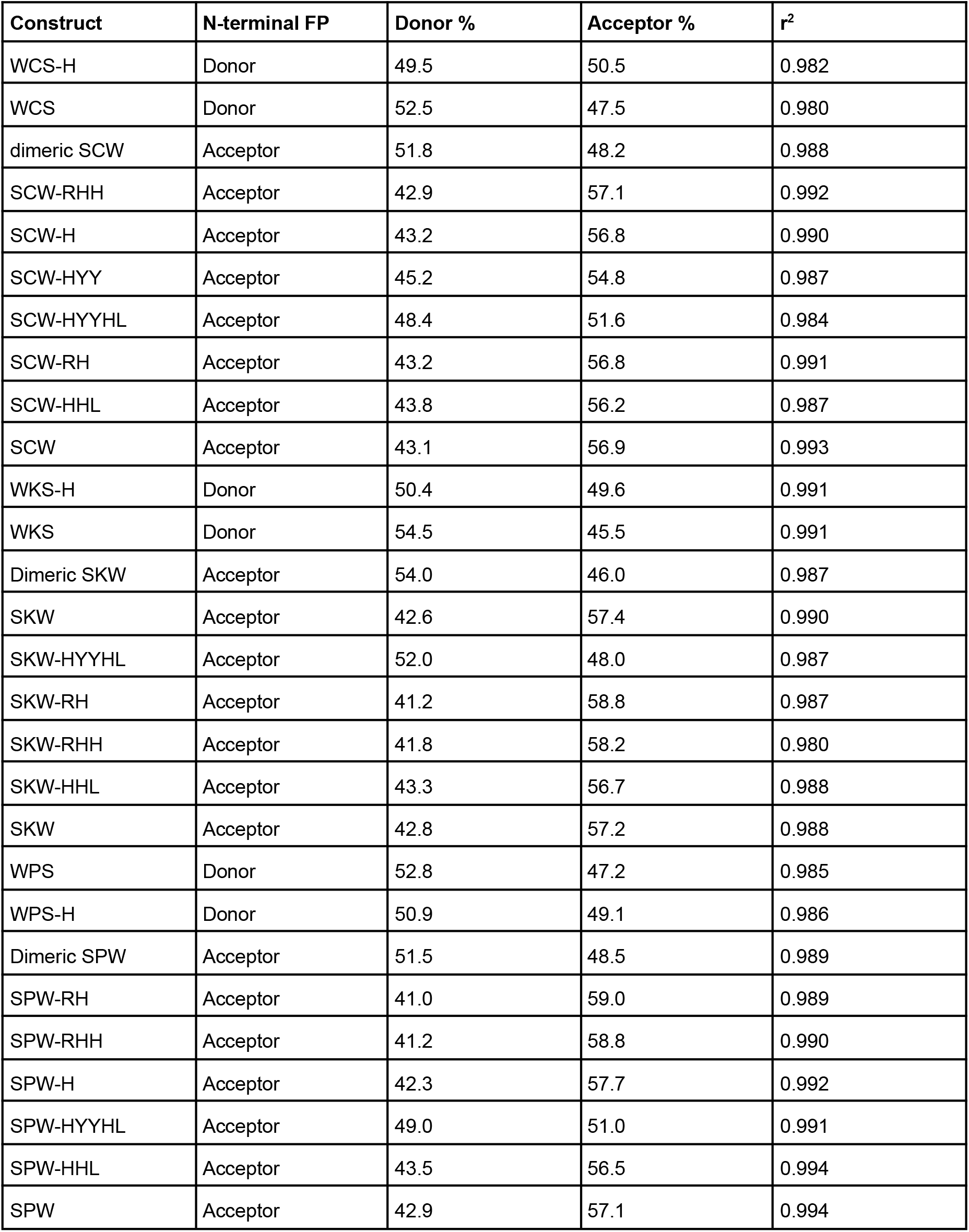
Percentages of acceptor and donor FP chromophores based on deconvoluted absorbance spectra of the FRET pairs.

| Construct | N-terminal FP | Donor % | Acceptor % | r<sup>2</sup> |
| --- | --- | --- | --- | --- |
| WCS-H | Donor | 49.5 | 50.5 | 0.982 |
| WCS | Donor | 52.5 | 47.5 | 0.980 |
| dimeric SCW | Acceptor | 51.8 | 48.2 | 0.988 |
| SCW-RHH | Acceptor | 42.9 | 57.1 | 0.992 |
| SCW-H | Acceptor | 43.2 | 56.8 | 0.990 |
| SCW-HYY | Acceptor | 45.2 | 54.8 | 0.987 |
| SCW-HYYHL | Acceptor | 48.4 | 51.6 | 0.984 |
| SCW-RH | Acceptor | 43.2 | 56.8 | 0.991 |
| SCW-HHL | Acceptor | 43.8 | 56.2 | 0.987 |
| SCW | Acceptor | 43.1 | 56.9 | 0.993 |
| WKS-H | Donor | 50.4 | 49.6 | 0.991 |
| WKS | Donor | 54.5 | 45.5 | 0.991 |
| Dimeric SKW | Acceptor | 54.0 | 46.0 | 0.987 |
| SKW | Acceptor | 42.6 | 57.4 | 0.990 |
| SKW-HYYHL | Acceptor | 52.0 | 48.0 | 0.987 |
| SKW-RH | Acceptor | 41.2 | 58.8 | 0.987 |
| SKW-RHH | Acceptor | 41.8 | 58.2 | 0.980 |
| SKW-HHL | Acceptor | 43.3 | 56.7 | 0.988 |
| SKW | Acceptor | 42.8 | 57.2 | 0.988 |
| WPS | Donor | 52.8 | 47.2 | 0.985 |
| WPS-H | Donor | 50.9 | 49.1 | 0.986 |
| Dimeric SPW | Acceptor | 51.5 | 48.5 | 0.989 |
| SPW-RH | Acceptor | 41.0 | 59.0 | 0.989 |
| SPW-RHH | Acceptor | 41.2 | 58.8 | 0.990 |
| SPW-H | Acceptor | 42.3 | 57.7 | 0.992 |
| SPW-HYYHL | Acceptor | 49.0 | 51.0 | 0.991 |
| SPW-HHL | Acceptor | 43.5 | 56.5 | 0.994 |
| SPW | Acceptor | 42.9 | 57.1 | 0.994 |

**Supplementary Table 3.**
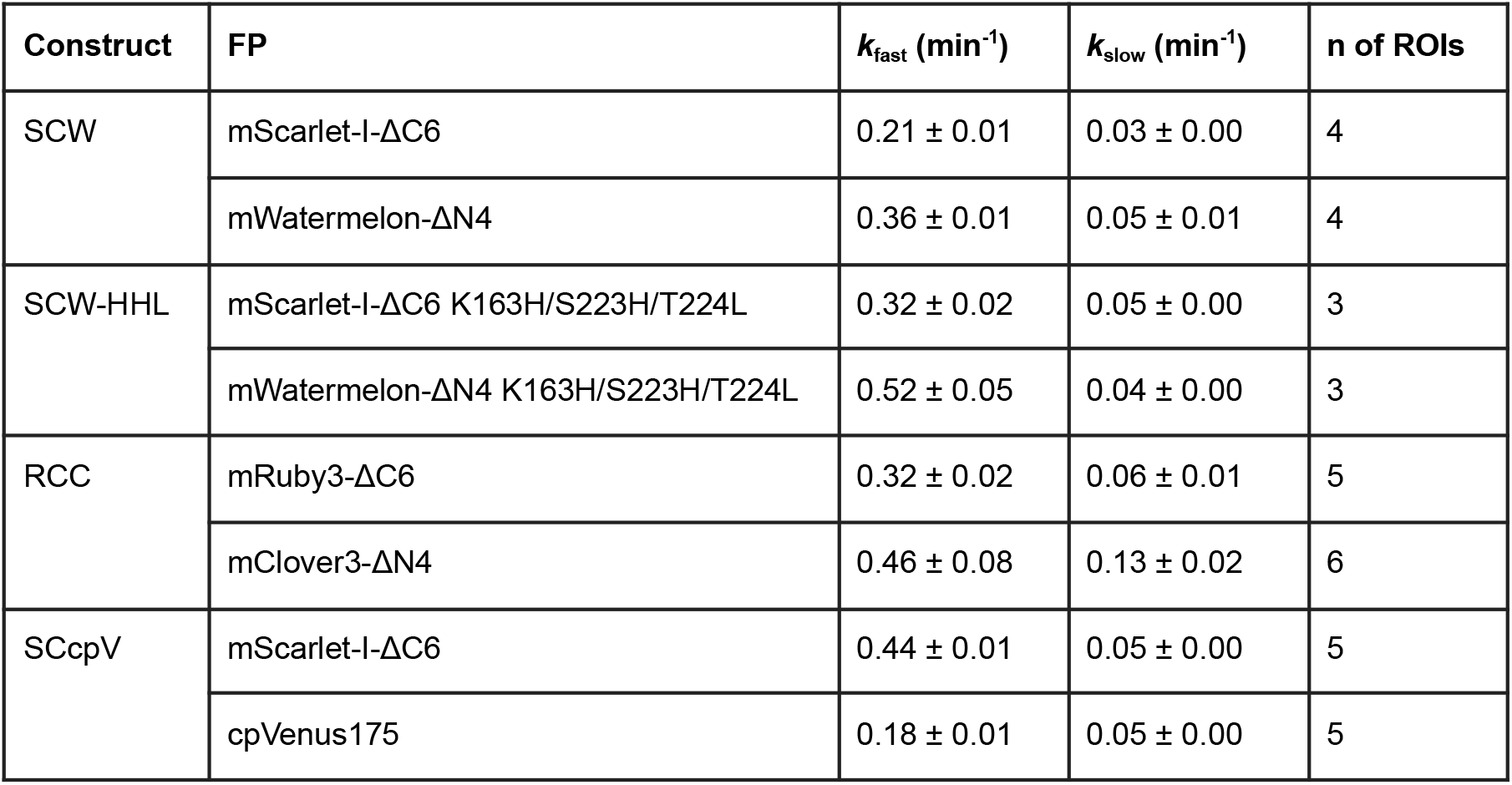
Photostability parameters of selected Ca^2+^ biosensors.

| Construct | FP | $k_{\text{fast}}$ (min<sup>-1</sup>) | $k_{\text{slow}}$ (min<sup>-1</sup>) | n of ROIs |
| --- | --- | --- | --- | --- |
| SCW | mScarlet-I-ΔC6 | 0.21 ± 0.01 | 0.03 ± 0.00 | 4 |
|  | mWatermelon-ΔN4 | 0.36 ± 0.01 | 0.05 ± 0.01 | 4 |
| SCW-HHL | mScarlet-I-ΔC6 K163H/S223H/T224L | 0.32 ± 0.02 | 0.05 ± 0.00 | 3 |
|  | mWatermelon-ΔN4 K163H/S223H/T224L | 0.52 ± 0.05 | 0.04 ± 0.00 | 3 |
| RCC | mRuby3-ΔC6 | 0.32 ± 0.02 | 0.06 ± 0.01 | 5 |
|  | mClover3-ΔN4 | 0.46 ± 0.08 | 0.13 ± 0.02 | 6 |
| SCcpV | mScarlet-I-ΔC6 | 0.44 ± 0.01 | 0.05 ± 0.00 | 5 |
|  | cpVenus175 | 0.18 ± 0.01 | 0.05 ± 0.00 | 5 |

**Supplementary Figure 1.**
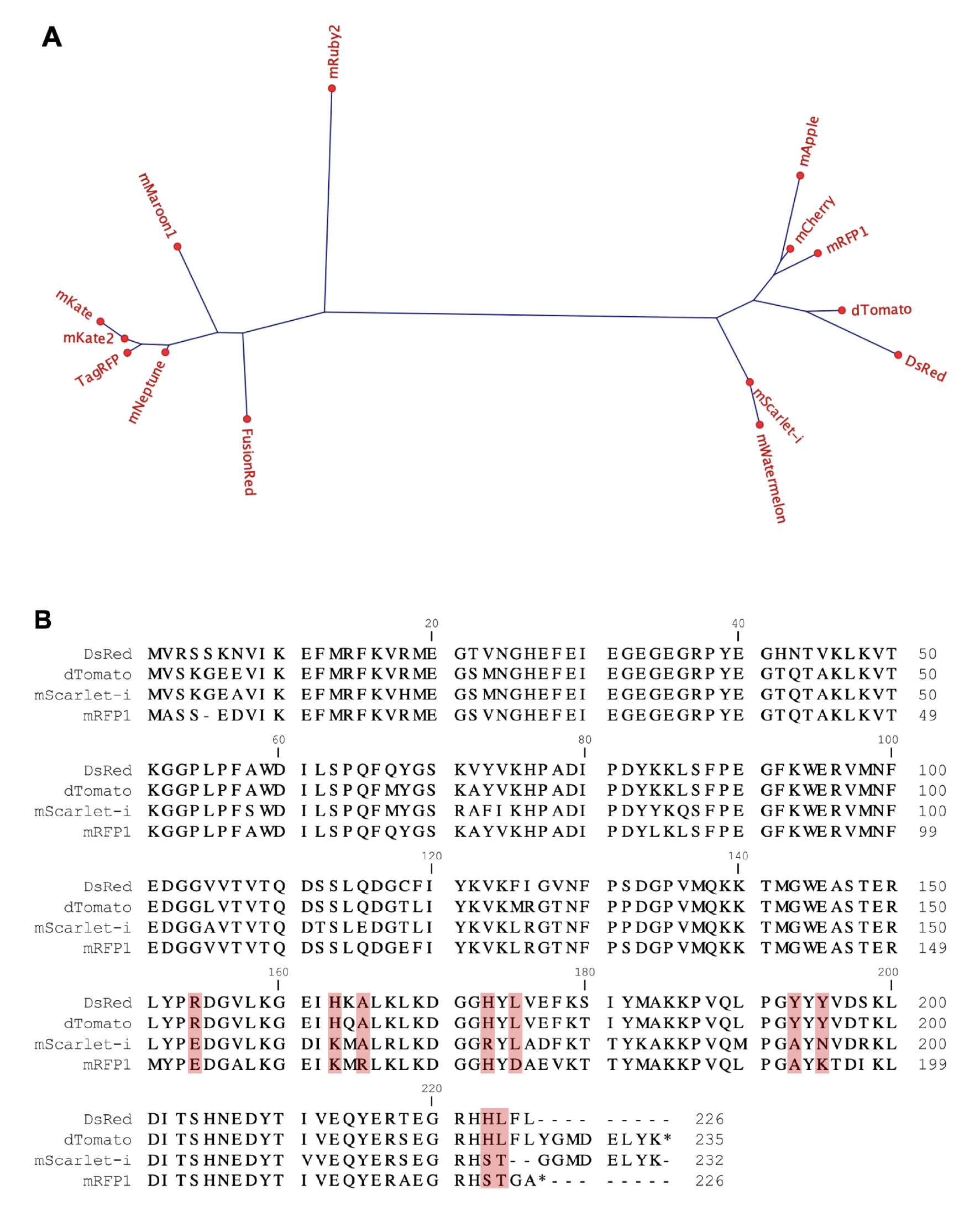
Sequence analysis of mScarlet-I. (A) Phylogenetic analysis of mScarlet-I, mWatermelon, and other RFPs. (B) Sequence alignment of mScarlet-I and DsRed-derived RFPs.

**Supplementary Figure 2.**
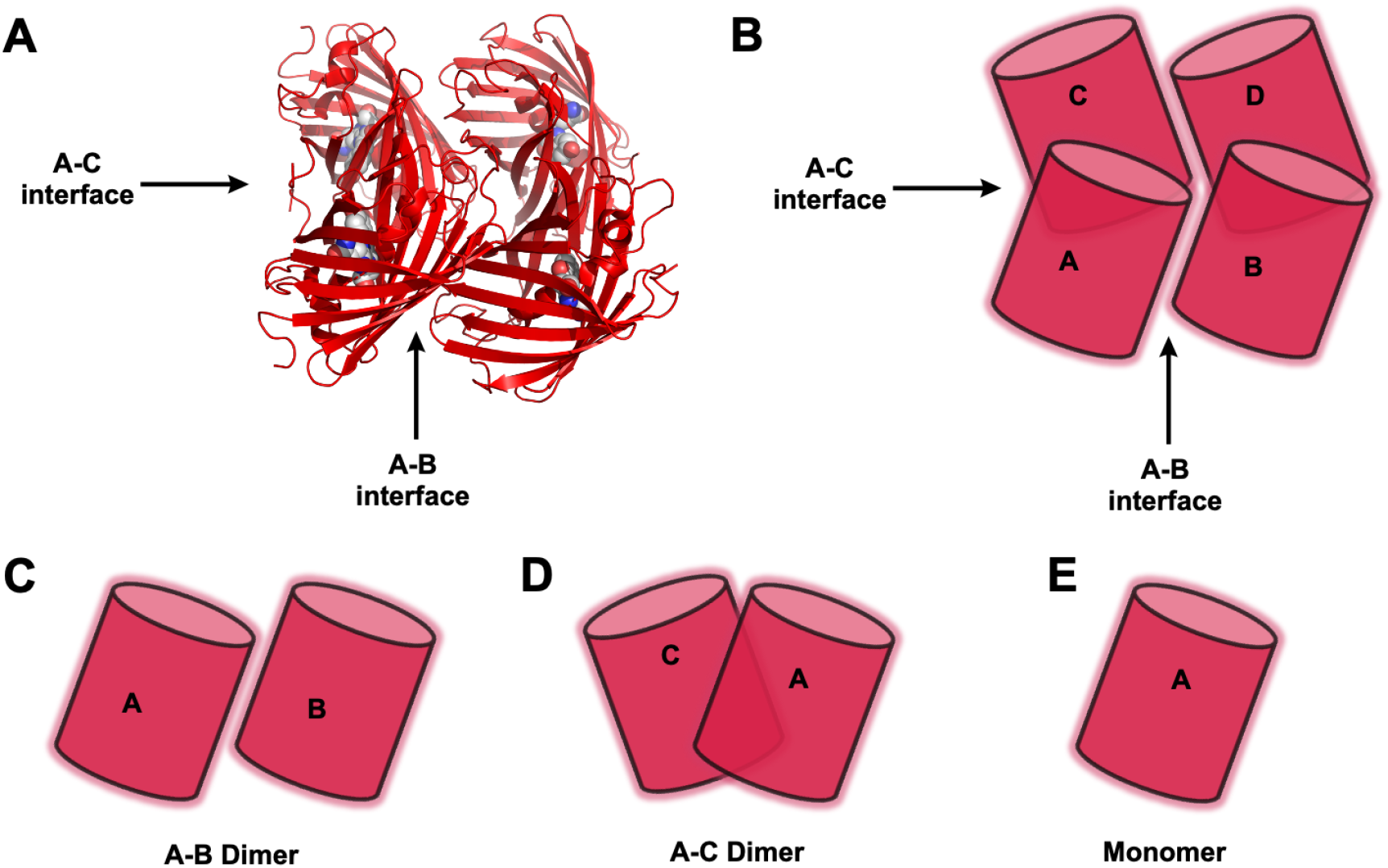
DsRed tetrameric structure.

**Supplementary Figure 3.**
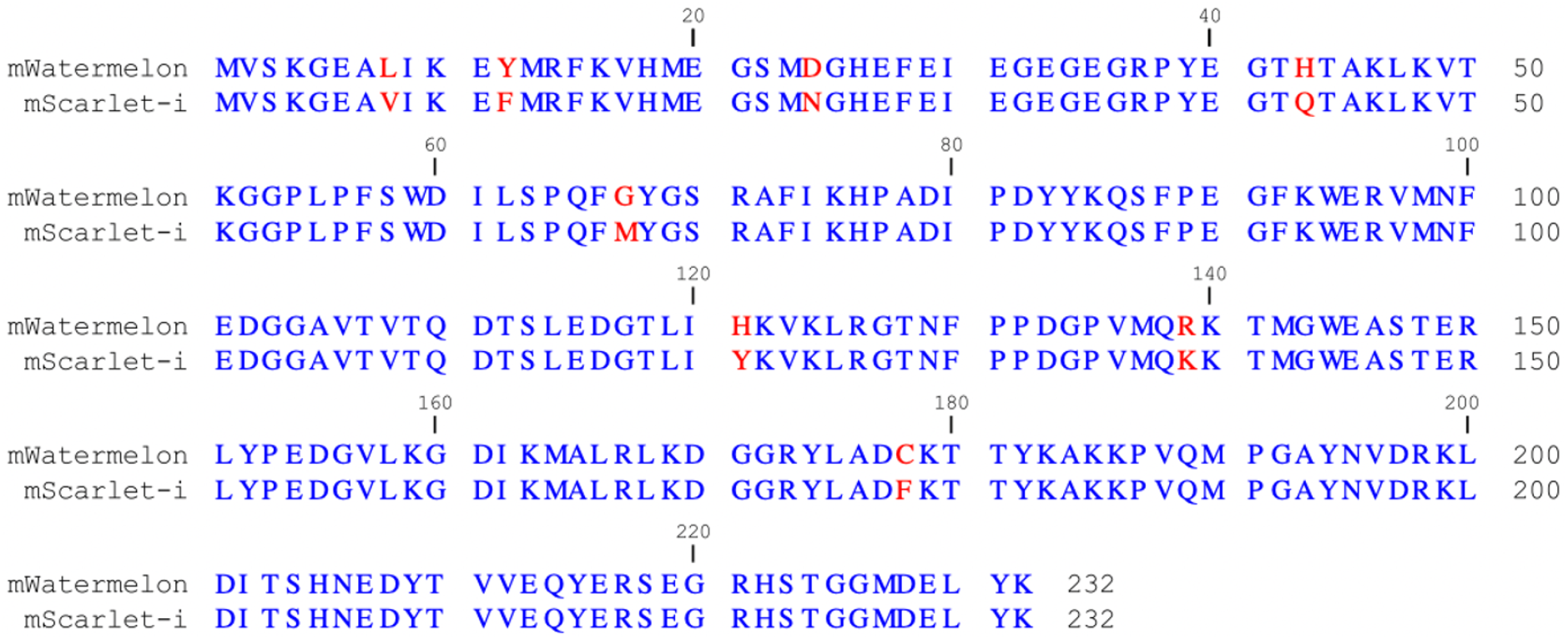
Sequence alignment for mWatermelon and mScarlet-I, mutation positions are highlighted in red.

**Supplementary Figure 4.**
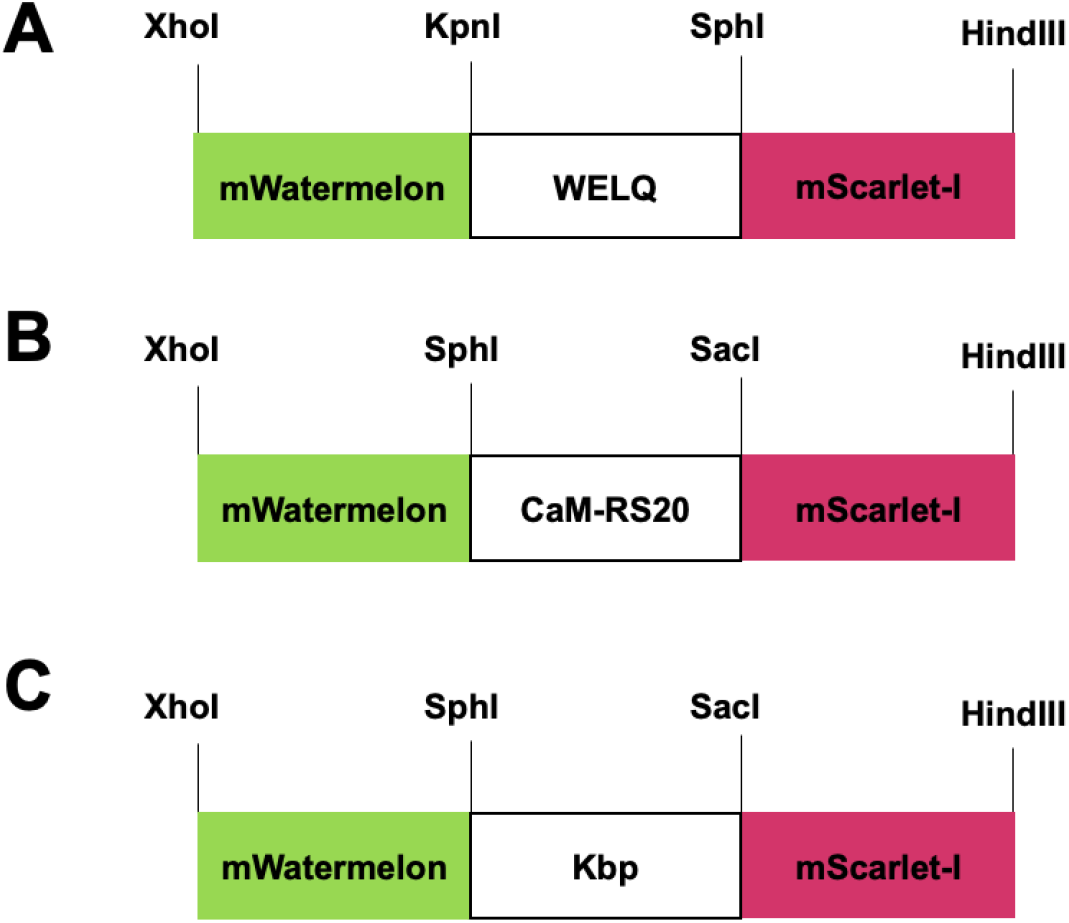
Design and restriction sites introduced in (A) protease biosensors, (B) Ca^2+^ biosensors, and (C) K^+^ biosensors.

**Supplementary Figure 5.**
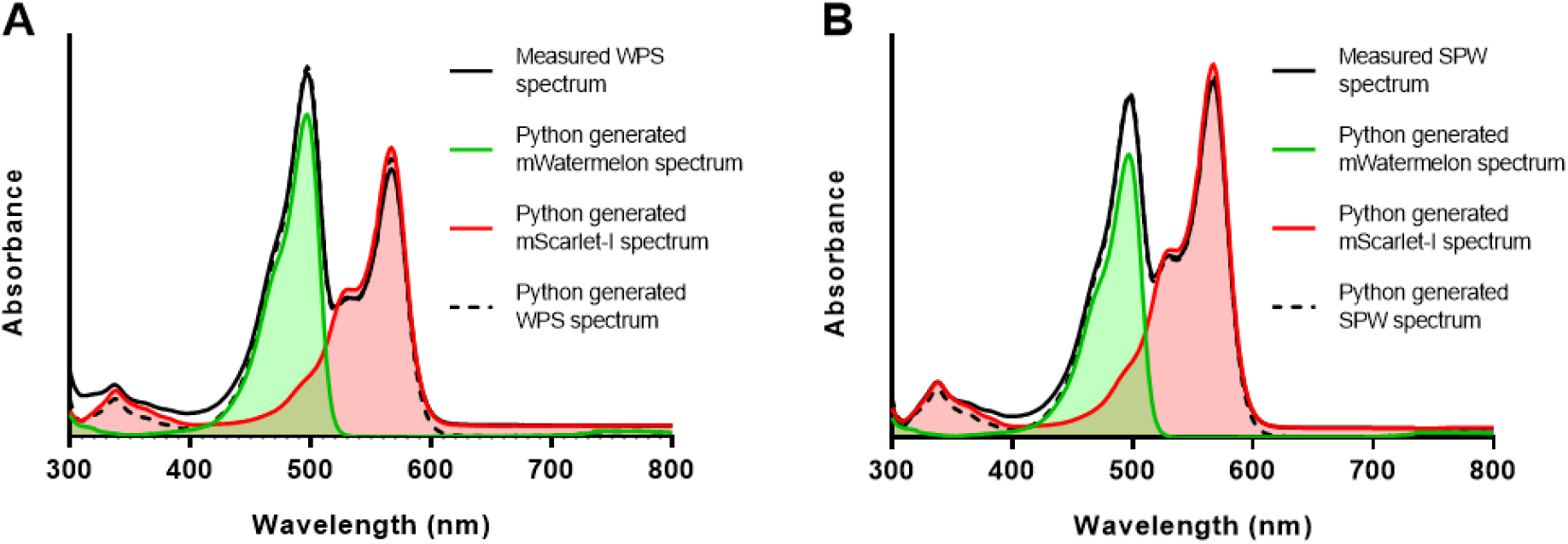
Deconvolution of absorbance spectra of (A) WPS and (B) SPW.

**Supplementary Figure 6.**
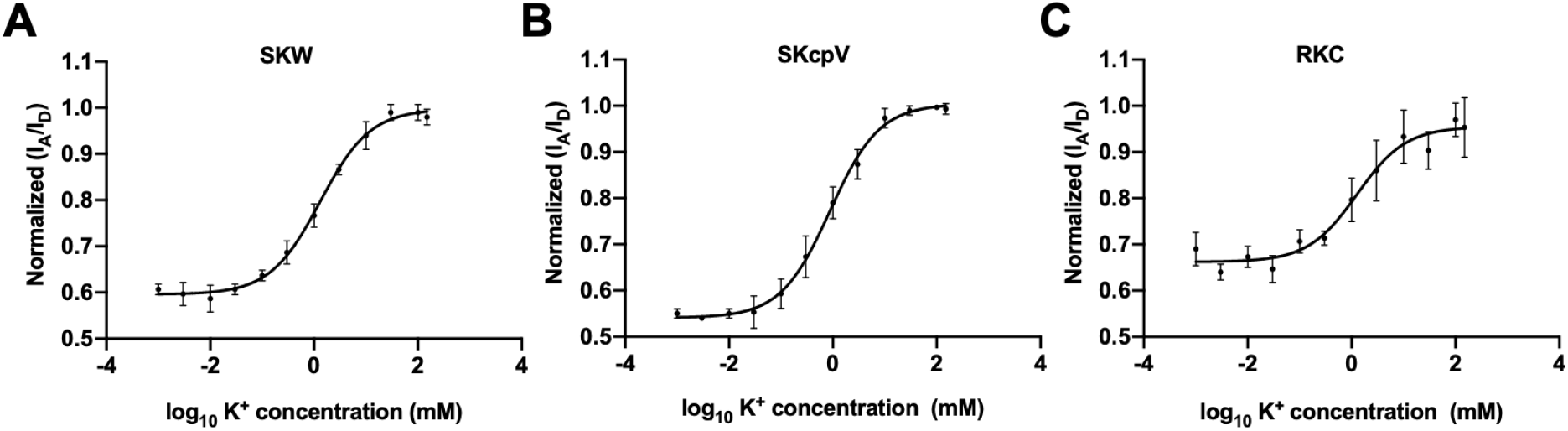
K^+^ titration curves of (A) SKW, (B) SKcpV, and (C) RKC are shown as normalized acceptor-to-donor fluorescence ratio versus log_10_ of K^+^ concentrations. Data are expressed as mean ± S.D for n = 3.

**Supplementary Figure 7.**
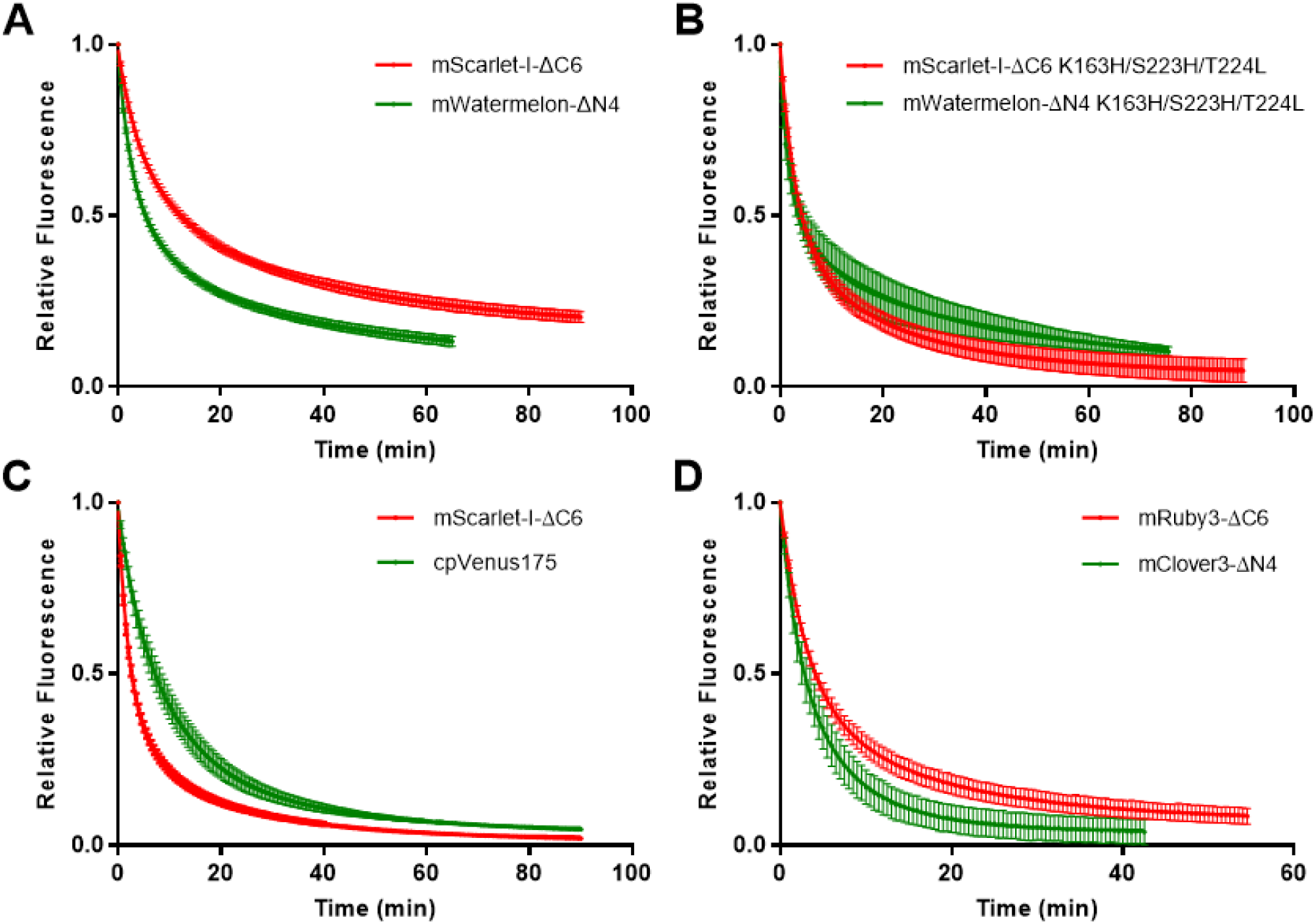
The fluorescence decay curves of selected FRET-based Ca^2+^ biosensors: (A) SCW, (B) SCW-HHL, (C) SCcpV, and (D) RCC. The curves are represented as mean ± S.D with the n (number of ROIs) listed in **Table S3**. Decay parameters *k*_fast_ and *k*_slow_ (**Table S3**) were obtained from the curve fitting with the two-phase decay model in Graphpad Prism.

